# Carriage of Supernumerary Sex Chromosomes Decreases the Volume and Alters the Shape of Limbic Structures

**DOI:** 10.1101/346767

**Authors:** Ajay Nadig, Paul K. Reardon, Jakob Seidlitz, Cassidy L. McDermott, Jonathan D. Blumenthal, Liv S. Clasen, Francois Lalonde, Jason P. Lerch, Mallar M. Chakravarty, Armin Raznahan

## Abstract

Sex chromosome aneuploidy (SCA) enhances risk for several psychiatric disorders associated with the limbic system, including mood and autism spectrum disorders. These patients provide a powerful genetics-first model for understanding the biological basis of psychopathology. Additionally, these disorders are frequently sex-biased in prevalence, further suggesting an etiological role for sex chromosomes. To clarify how limbic anatomy varies across sex and sex chromosome complement, we characterized amygdala and hippocampus structure in a uniquely large sample of patients carrying supernumerary sex chromosomes (n = 132) and typically developing controls (n=166). After correction for sex-differences in brain size, karyotypically normal males (XY) and females (XX) did not differ in volume or shape of either structure. In contrast, all SCAs were associated with lowered amygdala volume relative to gonadally-matched controls. This effect was robust to three different methods for total brain volume correction, including an allometric analysis that derived normative scaling rules for these structures in a separate, typically developing population (n = 79). Hippocampal volume was insensitive to SCA after correction for total brain volume. However, surface-based analysis revealed that SCA, regardless of specific karyotype, was consistently associated with a spatially specific pattern of shape change in both amygdala and hippocampus. In particular, SCA was accompanied by contraction around the basomedial nucleus of the amygdala and an area within the hippocampal surface that cuts across hippocampal subfields. These results demonstrate the power of SCA as a model to understand how copy number variation can precipitate changes in brain systems relevant to psychiatric disease.

## Significance Statement

In approximately 1 per 500 live births, infants are born carrying extra X- or Y-chromosomes. These conditions, known as sex chromosome aneuploidies (SCA), are associated with elevated risk for psychiatric disorders such as depression, anxiety, and autism. In our study, we leverage a uniquely large dataset of brain scans from SCA patients to characterize how SCA influences the structure of the amygdala and hippocampus, which have been strongly implicated in many SCA-linked neuropsychiatric disorders. Across our different patient groups, we find converging evidence that SCA is associated with a decrease in amygdala volume, and spatially specific areal contractions in both the amygdala and hippocampus. Our findings help clarify how genetic copy number variation associated with psychiatric disease shapes brain organization.

## Introduction

The amygdala and hippocampus are key elements of a limbic brain network that is perturbed in diverse neuropsychiatric disorders, including mood disorders (Price & Drevets, 2012) and autism (Hennessey et al. 2018; Amaral et al. 2008). Patients with sex chromosome aneuploidies (SCA) are at elevated risk for these psychiatric disorders (Hong & Reiss, 2014), although it remains unclear if amygdalar and hippocampal development are sensitive to the presence of supernumerary sex chromosomes. Some evidence that the volume and shape of these structures may be sensitive to SCA comes from recent studies of Turner Syndrome which report that loss of one X-chromosome in females enlarges the amygdala in a spatially specific manner (Green et al., 2016; Kesler et al., 2004). However, it remains unknown if these effects generalize to the hippocampus (the other core element of the limbic brain network), or if either of these structures are sensitive to carriage of excess X- and or Y-chromosomes. There are no existent studies of Y-chromosome dosage effects on amygdala and hippocampus structure. Clarifying the nature of SCA effects on amygdalar and hippocampal anatomy is not only relevant for the neurobiology of sex chromosome aneuploidies, but also informs the broader question of sex-differences in amygdalar and hippocampal anatomy. The fact that many disorders linked to limbic dysfunction also show clear sex-biases in prevalence (anxiety, 1.8x risk for females versus males: Ruscio et al. 2017; autism, 4x risk for males versus females: Werling and Geschwind 2013) could potentially reflect sensitivity of limbic structures to biological factors that differ between males and females, such as X- and Y-chromosome dosage.

Here, we detail sex and SCA effects on anatomy of the amygdala and hippocampus by applying recently developed methods for morphometric analysis (Chakravarty et al., 2013) to a large set of structural MRI scans of individuals with varying sex chromosome dosage (n=298: 87 XX, 79 XY, 28 XXX, 56 XXY, 25 XYY, 19 XXYY, 5 XXXXY). A key methodological consideration in pursuing this goal is that sex chromosome aneuploidies are known to induce significant shifts in total brain volume (Raznahan et al., 2016), which may bias estimation of regional volumes given known patterns of non-linear subcortical scaling (Reardon et al. 2016; Reardon et al. 2018). Therefore, we use a separate, typically developing sample (N = 79) to derive normative allometric scaling rules for the amygdala and hippocampus, and assess patient/control volume contrasts in the context of these scaling norms. Lastly, we determine whether changes in volume are accompanied by changes in the shape of limbic structures by mapping sex and SCA effects on local surface area across 5245 points (vertices) across the amygdalo-hippocampal surface. Moving beyond analysis of bulk volume to study shape is critical given the complex topography of functional and connectional gradients within the amygdala and hippocampus (Carr, Rissman, & Wagner, 2010; Janak & Tye, 2015; Saygin et al., 2015).

## Method

### Sample

Our core sample included 298 individuals of varying karyotype. Our allometric sample, used to generate scaling rules for the amygdala and hippocampus, included 79 typically developing XX and XY individuals. Sample characteristics are described in Table 1. Participants with SCA were recruited through the National Institutes of Health (NIH) website and parent support groups; X-/Y-supernumeracy was confirmed by karyotype test. Non-mosaicism was confirmed by visualization of 50 metaphase spreads in peripheral blood. We excluded participants with a history of head injury, or gross brain abnormality. Typically developing individuals were drawn from the NIH Longitudinal Structural MRI Study of Human Brain Development (Giedd et al., 2015). Exclusion criteria for these participants included use of psychiatric medication, enrollment in special education services, history of mental health treatment, or prior diagnosis of a nervous system disorder.

**Table 1.** Participant characteristics.

| Group | XX | XY | XXX | XXY | XYY | XXYY | XXXXY | Allometric Sample |
| --- | --- | --- | --- | --- | --- | --- | --- | --- |
| *n* | 87 | 79 | 28 | 56 | 25 | 19 | 5 | 79 (34 F) |
| **Age (years)** |  |  |  |  |  |  |  |  |
| Mean (SD) | 12.7 (5.1) | 12.7 (4.6) | 12.3 (5.7) | 12.7 (4.9) | 12.2 (4.9) | 14.0 (5.5) | 12.9 (4.8) | 13.0 (0.6) |
| Range | 5.4-25.1 | 5.2-25.5 | 5.0-24.8 | 5.2-26.0 | 5.7-23.0 | 5.0-23.0 | 7.7-17.2 | 12.0 – 14.0 |
| **FSIQ** |  |  |  |  |  |  |  |  |
| Mean (SD)* | 113.9 (12.3) | 116.6 (13.3) | 94.1 (14.2) | 97.6 (17.1) | 91.0 (14.6) | 86.9 (12.9) | 55.7 (7.23) | 116.3 (12.4) |
| **SES** |  |  |  |  |  |  |  |  |
| Mean (SD)* | 47.3 (17.1) | 48.0 (21.3) | 40.9 (15.9) | 56.0 (20.7) | 58.1 (22.4) | 45.5 (22.6) | 68.8 (22.4) | 34.1 (19.7) |

FSIQ, Full Scale IQ; SES, socioeconomic status. \*p < 0.01 for omnibus test of significant variation across groups in core sample.

### Image acquisition and processing

We collected T1-weighted MRI scans for each individual on the same 1.5 Tesla General Electric SIGNA scanner with contiguous 1.5 mm axial slices using a 3D spoiled gradient-recalled echo sequence with the following acquisition parameters: echo time, 5 ms; repetition time, 24 ms; flip angle, 45°; acquisition matrix, 256 × 192; number of excitations, 1; field of view, 24 cm.

We estimated total brain volume by submitting scans to the CIVET 1.1.10 pipeline for automated morphometric analysis (Ad-Dab’bagh et al., 2006). This total brain volume estimate was the sum of total grey and white matter volume estimates (mm^3^) which were computed by CIVET tissue segmentation that used a validated neural net approach to voxel classification (Cocosco, Zijdenbos, & Evans, 2003; Zijdenbos, Forghani, & Evans, 2002).

All amygdala and hippocampus measurements were automatically generated by a well-validated multi-atlas segmentation algorithm, MAGeT Brain (Chakravarty et al., 2013; Pipitone et al., 2014). Briefly, this algorithm makes use of high-resolution in-vivo atlases generated high resolution T1 and T2 weighted images (3T, final isotropic voxel dimension 0.3 mm) from 5 reference subjects (3 male, 2 female) (Winterburn et al. 2013) that are aligned to 21 randomly selected participants within the NIH Human Brain Development in Health Study (Giedd et al., 2015), as has been previously described (Reardon et al. 2018). This pipeline produces a study-specific library of 105 amygdala and hippocampus segmentations (5 atlases × 21 templates). The final segmentation is decided upon using a voxel majority vote, where the most frequently applied label at each location is retained. These methods are reliable (Dice kappa = 0.86). A quality control image file is provided for each scan, allowing detailed visual inspection to rule out gross segmentation errors. All scans included in this analysis passed quality control of raw scans and MAGeT Brain segmentations for motion artifacts and other confounders by a rater who was blinded to subject karyotype (P.K.R.)

To determine shape, surface-based representation of amygdala and hippocampus structures were estimated using the marching cubes algorithm and morphologically smoothed using the AMIRA software package (Visage Imaging). Next, the nonlinear portions of the 21 transformations mapping subjects to the 21 input templates were concatenated and averaged across input templates to limit the effects of noise and error. These surface based representations were warped to fit each template, and each surface was warped to match each subject. This procedure yields 30 possible surface representations per subject that are then merged by estimating the median coordinate representation at each location. At this point, one-third of the surface of each triangle is assigned to each vertex within the triangle. The surface area value stored at each vertex is the sum of all such assignments form all connected triangles. Lastly, surface-area values were blurred with a surface based diffusion-smoothing kernel (5 mm). The final derived measures of interest were total volume estimates for the left and right amygdala and hippocampus, and surface area at 5245 vertices across all structures (right amygdala: 1405; left amygdala: 1473; right hippocampus: 1215; left hippocampus: 1152).

### Statistical Analyses

#### Bulk volume comparisons

We first quantified the effect of karyotype on total bilateral amygdala and hippocampus volume in the core sample using omnibus *F* tests. For structures showing a significant omnibus effect of group, follow-up *post hoc* pairwise *t* tests were used to specify pairwise karyotype group differences in structure volume. To simultaneously visualize changes in regional and total brain volumes across karyotype groups, we plotted volumetric differences as an effect-size shift relative to the distribution observed in XY males. Effect sizes were calculated as (mean volume of XY group - mean volume of comparison group)/SD of XY group.

#### Normative allometry of the amygdala and hippocampus

We used a classic log-log regression approach (Huxley, 1924) to derive normative scaling rules for the amygdala and hippocampus within the independent allometric sample of 79 typically developing males and females. The scaling relationship between a regional volume and total brain volume is given by the β_1_ coefficient in the following equation:

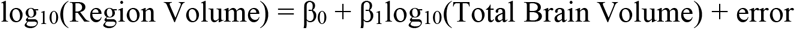

This method allows simple quantification of non-linear allometric scaling relationships between regional and total brain volume. Furthermore, the scaling coefficient from this fit is easily interpretable: β_1_ values significantly less than 1 indicate a hypoallometric scaling relationship (i.e. regional volume becomes proportionally smaller as total brain volume becomes larger); β_1_ values significantly greater than 1 indicate a hyperallometric scaling relationship (i.e. regional volume becomes proportionally larger as total brain volume becomes larger); β_1_ values equal to 1 indicate an isometric scaling relationship (proportional size of the region is maintained across the range of total brain volume).

To account for sex-differences in the allometric fit β_0_ and β_1_ that have been observed in other brain structures (Mankiw et al., 2017; Reardon et al., 2016), we used our log-log regression framework to sequentially test three general linear models of amygdala and hippocampus allometry with decreasing complexity of sex differences.

First, we tested for sex differences in the scaling coefficient itself, such that different allometric scaling principles hold for males and females (β_3_):

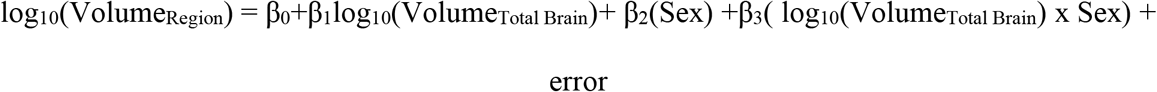

In the absence of evidence for a sex difference in scaling (p>0.05), we proceeded to the next model, which includes a single scaling relationship that allows for baseline sex difference in regional volume, reflected in the β_2_term:

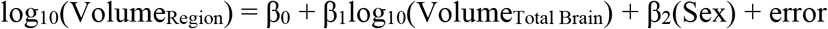

In the absence of evidence for a sex difference in volume (p>0.05), we proceeded to the last model, the simple allometric model with no sex term:

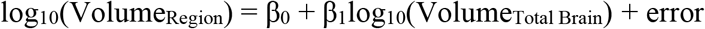

#### Sex and SCA effects in the context of allometry

We used the scaling rules derived above to test for brain size-independent effects of sex chromosome complement on amygdala and hippocampus volumes. For each SCA group in our core sample (excluding the small sample of five XXXXY individuals), we tested for differences between observed regional volumes and regional volumes predicted from total brain volume. For each regional volume, we selected the appropriate allometric model using the procedure defined above, and predicted regional volume for each individual in the core sample based on total brain volume and sex (including sex only if a significant sex difference in normative scaling was observed). We then calculated the deviation of each individual’s observed log_10_(Volume_Region_) from the predicted log_10_(Volume_Region_) with simple subtraction. Finally, we compared the distribution of these residuals between each SCA group and its appropriate gonadal control within the core sample (i.e. XXX vs XX, and other SCAs vs. XY) using a regression approach with the following formula:

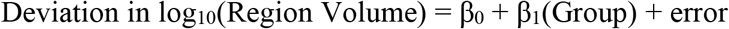

where Group was a binary categorical variable. For each region, *p* values from these tests were Bonferroni corrected to account for the five SCA group contrasts being performed.

#### Analysis of sex and SCA effects using normalization and covariation to control for brain size

We also compared the results of our allometric approach with two complementary approaches that are commonly used in the neuroanatomical literature: normalization and covariation. In normalization, group differences are tested after re-expressing regional volume as a fraction of total brain volume. In covariation, group differences are tested with total brain volume included as a covariate.

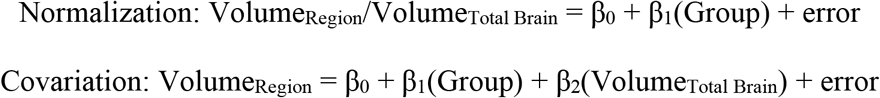

To assess SCA effects, we ran these two models for each SCA group and its respective gonadal control group, such that Group was a binary categorical variable. Similar to our allometric approach, we Bonferroni corrected *p* values for each region to account for the five contrasts being performed. To assess sex effects, we ran these models in our allometric sample, with Group reflecting sex.

All statistical analyses described above were performed using R software (R Core Team, 2008). Data were visualized using the *ggplot2* (Wickham, 2016) and *patchwork* (Pendersen, 2017) packages.

#### Vertex-wise normative allometry of the amygdala and hippocampus surface

In order to assess if carriage of supernumerary sex chromosomes distorts amygdalar and hippocampal shape, we developed normative models for the relationship between shape and size of these structures. Because the MAGeT Brain morphometric pipeline represents structure shape as vertex-wise measures of local surface area, relationships between structure size and shape can be captured by modelling inter-individual variation in area at each vertex by inter-individual in total structure surface area, which is strongly related to total structure volume (Pearson’s correlation between total region volume and total region surface areas: 0.97, amygdala; 0.89, hippocampus). We used a tiered approach to modelling sex differences similar to our bulk volume analyses. Critically, although vertex-wise variation in significance of sex differences is possible, for the purposes of simplicity and comparability of model parameters across vertices, we required that all vertices use the same final model. Thus, in our normative allometric sample (n = 79), we first modelled each vertex surface area estimate using the following equation:

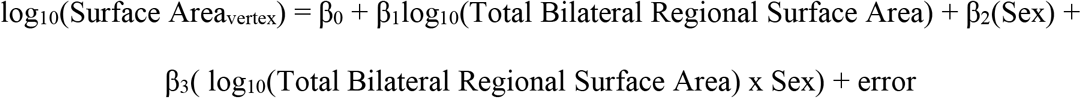

If any vertices contained significant interaction terms (p_FDR_<0.05), we applied this model to every vertex. In the absence of any vertices containing significant interaction terms, we proceeded to the next lower order vertex-wise model:

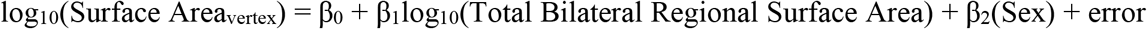

If any vertices contained significant sex terms (p_FDR_<0.05), we applied this model to every vertex. Finally, in the absence of any vertices containing significant sex terms, we proceeded to the lowest order, simple allometric vertex-wise model:

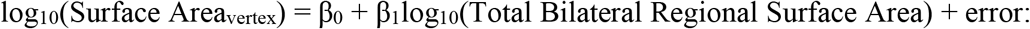

Vertex-wise β_1_ coefficients were visualized to represent which subregions scale hypo-, iso-, or hyper-allometrically with total regional surface area.

### Vertex-wise changes in surface area across SCA in the context of normative surface allometry

We performed a similar correction procedure as was applied in our volumetric analysis to assess allometrically-corrected changes in surface area across karyotype groups. For each vertex, we predicted log_10_(Surface_vertex_), for each individual in the core sample based on total regional surface area and sex. We then calculated the deviation of each individual’s actual log_10_(Surface_vertex_) from the predicted log_10_(Surface_vertex_) at each vertex with simple subtraction. To assess SCA, we applied a simple model at each vertex separately for each contrast:

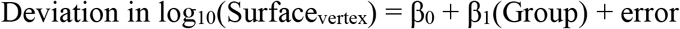

where Group was a binary categorical variable. We corrected *p* values separately for each contrast, for each region, using the False Discovery Rate with q (expected proportion of falsely - rejected nulls) set at 0.05.

## Results

### Total brain volume, raw amygdala volume, and raw hippocampus volume are patterned across sex and sex chromosome complement

Total brain volume, amygdala volume, and hippocampus volume showed statistically significant variation across the seven distinct karyotype groups represented in our core sample of 298 individuals (Table 2, Fig. 1A). XX females had significantly smaller total brain, raw amygdala, and raw hippocampus volumes than XY males. Analysis of SCA revealed that carriage of a supernumerary X-chromosome was associated with decreased amygdala volume compared to gonadal controls. These differences were statistically-significant for all SCAs, and increased in effect size with mounting sex chromosome dosage. Carriage of a supernumerary X-chromosome was associated with decreased hippocampus volume compared to gonadal matched controls, except in the case of XXYY. In the small (n=19) XXYY karyotype, we observed a non-significant reduction in hippocampus volume that was trend-level before correction for multiple comparisons (*p*_uncorrected_= 0.08), although this karyotype group was associated with the greatest effect size reduction in hippocampal volume. The XYY karyotype was not associated with significant differences in total brain volume, raw amygdala volume, or raw hippocampus volume as compared to XY males.

**Figure 1.**
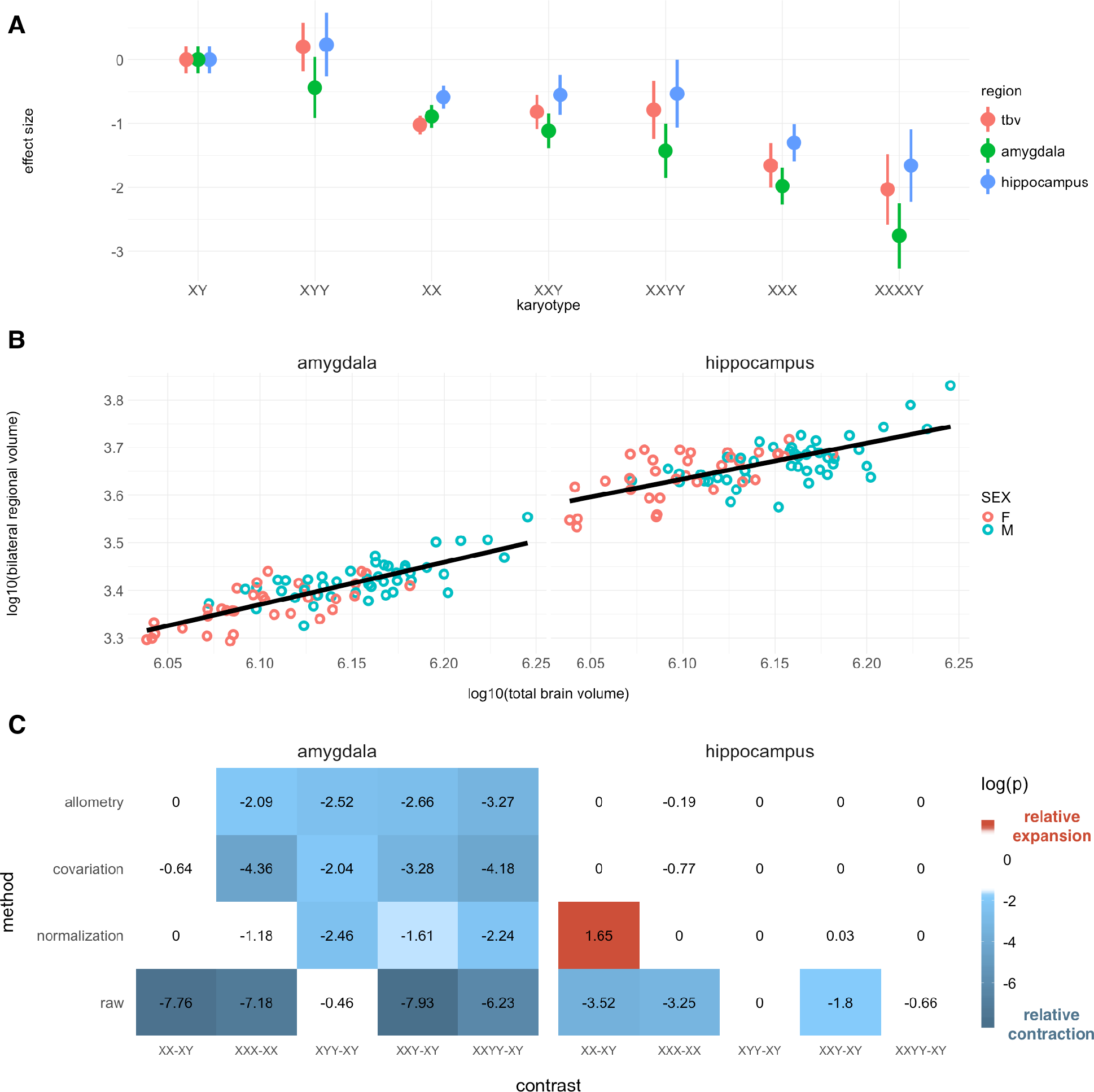
Bulk volume analysis of amygdala and hippocampus. (A) Total brain volume (tbv), amygdala, and hippocampus volume across karyotype groups, relative to typically developing males (XY). (B) Scaling of amygdala and hippocampus volume in the allometric sample (amygdala β_1_ = 0.89; hippocampus β_1_ = 0.76). (C) Comparisons of regional volume across sex and sex chromosome complement. FDR-corrected log(p) values are shown. Presence of color indicates statistical significance, and color identity/sign indicates direction of effect: for contrast A-B, blue and negative values indicate smaller volume in A versus B, red and positive value indicate larger volume in A versus B.

**Table 2.**
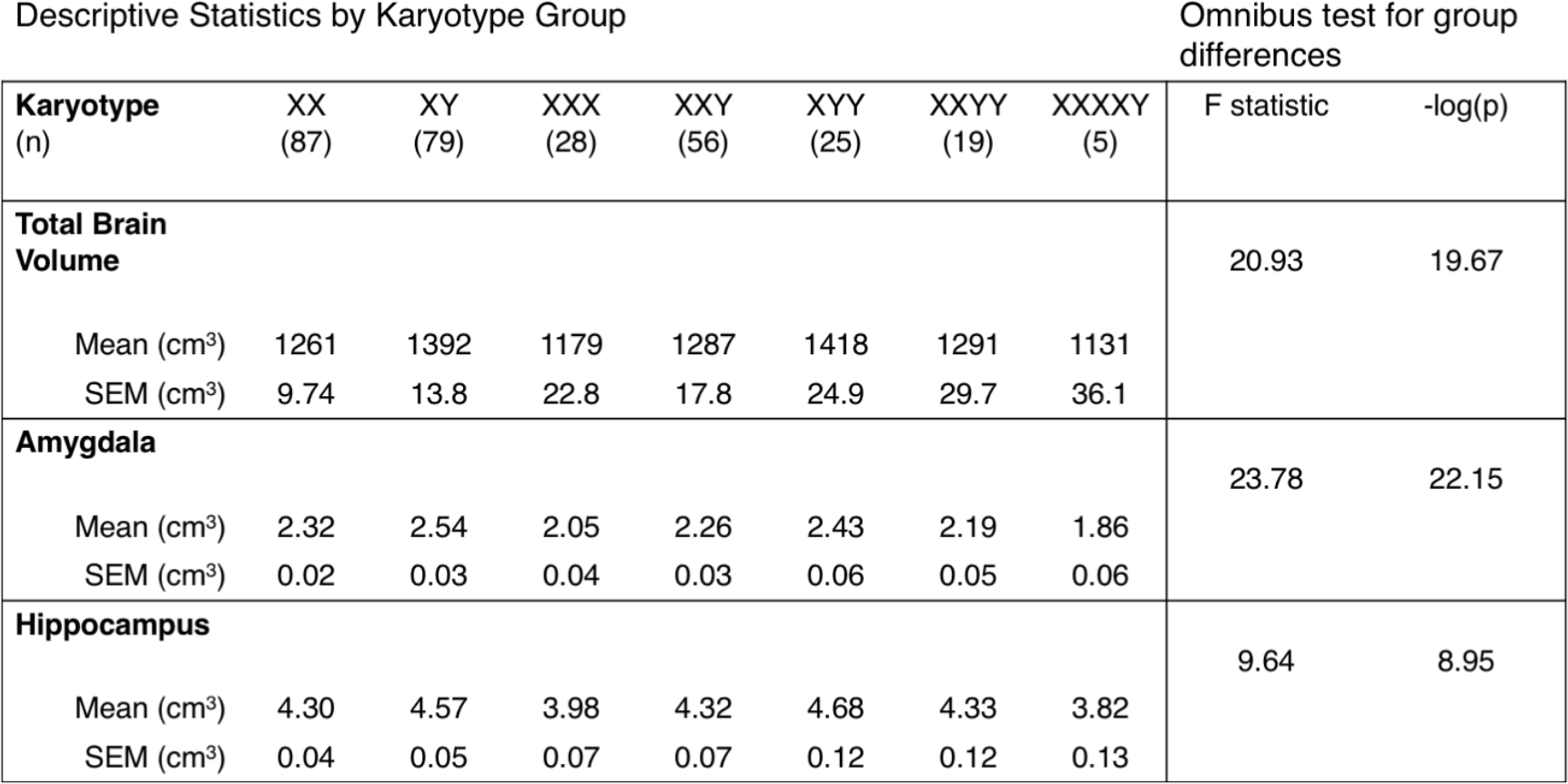
Descriptive statistics for bulk volume analysis.

| Karyotype (n) | XX (87) | XY (79) | XXX (28) | XXY (56) | XYY (25) | XXYY (19) | XXXXY (5) | F statistic | -log(p) |
| --- | --- | --- | --- | --- | --- | --- | --- | --- | --- |
| **Total Brain Volume** |  |  |  |  |  |  |  | 20.93 | 19.67 |
| Mean (cm³) | 1261 | 1392 | 1179 | 1287 | 1418 | 1291 | 1131 |  |  |
| SEM (cm³) | 9.74 | 13.8 | 22.8 | 17.8 | 24.9 | 29.7 | 36.1 |  |  |
| **Amygdala** |  |  |  |  |  |  |  | 23.78 | 22.15 |
| Mean (cm³) | 2.32 | 2.54 | 2.05 | 2.26 | 2.43 | 2.19 | 1.86 |  |  |
| SEM (cm³) | 0.02 | 0.03 | 0.04 | 0.03 | 0.06 | 0.05 | 0.06 |  |  |
| **Hippocampus** |  |  |  |  |  |  |  | 9.64 | 8.95 |
| Mean (cm³) | 4.30 | 4.57 | 3.98 | 4.32 | 4.68 | 4.33 | 3.82 |  |  |
| SEM (cm³) | 0.04 | 0.05 | 0.07 | 0.07 | 0.12 | 0.12 | 0.13 |  |  |

### The amygdala and hippocampus scale hypoallometrically with total brain volume

In our allometric sample of 79 typically developing individuals, there were no significant effects of sex on either the intercepts or slopes of allometric models for amygdala or hippocampal volume (see Methods), so we proceeded with the simplest allometric model: log_10_(Volume_Region_) = β_0_ + β_1_log_10_(Volume_Total Brain_). Both amygdala and hippocampus volume scaled hypoallometrically with total brain volume (Fig.1B, Amygdala: β_1_=0.89; Hippocampus: β_1_= 0.76).

### A subset of sex and sex chromosome complement effects on amygdala and hippocampus volume survive allometric correction

To assess whether changes in raw amygdala and hippocampus volume as a function of sex and SCA represent distortions of normative brain proportionality, we examined deviations from derived normative scaling rules across sex and sex chromosome complement (Fig. 1C). Sex differences in amygdala and hippocampus volume were not statistically-significant after allometric correction for total brain volume. The decrease in amygdala volume with carriage of a supernumerary X-chromosome remained significant after allometric correction. Allometric correction revealed a statistically-significant amygdala volume decrease in XYY versus XY males, where no significant change was detected in raw volumes. These results indicate a convergent effect of supernumerary X- and Y-chromosome carriage on proportional amygdala volume. Differences in hippocampus volume across sex and sex chromosome complement were eliminated by allometric correction, indicating that proportional hippocampus volume is maintained across karyotype groups.

Results from normalization and covariation analysis were largely consistent with those of allometric analysis, with two notable exceptions. A significant decrease of hippocampus volume in males versus females was detected by normalization analysis, where an opposite effect was detected in raw volumes, and no significant effect was detected by covariation or allometric analysis. Also, decrease in amygdala volume in XXX versus XX groups was not significant in normalization analysis, but was significant in all other comparison techniques.

### Vertex-wise normative allometry of the amygdala and hippocampus

To assess how sex and sex chromosome complement impact regional shape, we first developed a normative model for scaling of amygdala and hippocampus shape with size in our allometric sample. We found no significant interactive or additive effects of sex in our allometric model, leading us to use the simplest vertex-wise log-log regression between vertex surface area and total bilateral region surface area. We found that as total bilateral surface area of the amygdala and hippocampus increased, subregions scaled heterogeneously (Fig. 2A). For the amygdala, the rostral medial surface scaled hyperallometrically, whereas the caudal lateral surface scaled hypoallometrically. For the hippocampus, the rostral and lateral surfaces scaled hyperallometrically, whereas the medial caudal surface scaled hypoallometrically.

**Figure 2.**
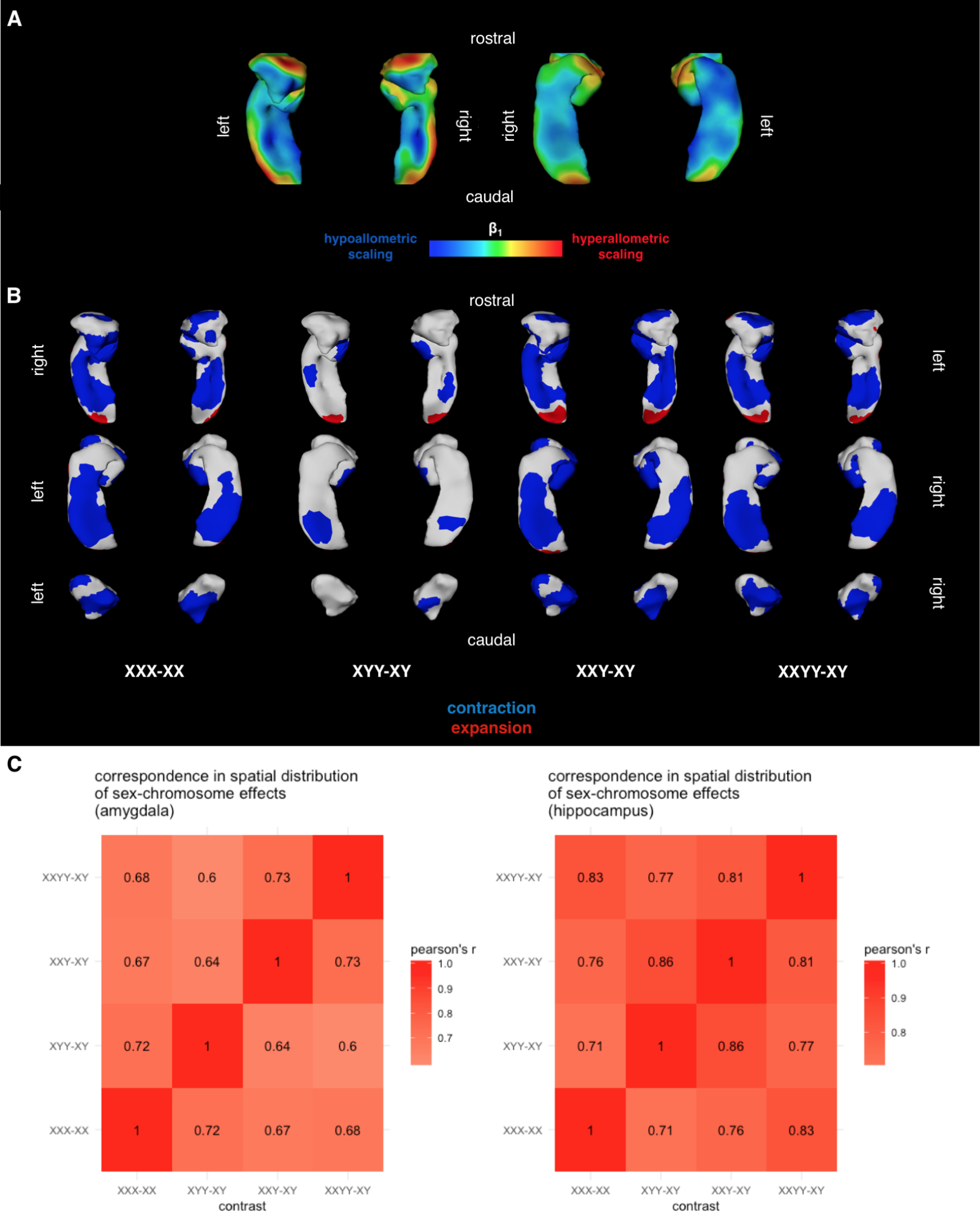
Surface-based analysis of amygdala and hippocampus shape. (A) Map of vertex-wise surface area scaling coefficients from log-log linear model with total bilateral region surface area. (B) Binarized surface maps showing areas of significant surface area contraction (blue) or expansion (red) for each patient contrast. (C) Spatial correlation of uncorrected t-statistic maps demonstrating convergence of SCA effects.

### Amygdala shape is systematically altered by SCA

Our normative description of amygdala scaling in health provided a foundation to assess whether amygdala shape is atypical in SCA. For each participant’s scan, we computed vertex-wise deviations in surface area from normative scaling predictions based on total bilateral amygdala surface area, and asked whether these deviations differed across the karyotypic contrasts of interest.

After FDR correction, we observed statistically significant changes in proportional surface area in each patient group (Fig 2B). In particular, we identified convergent changes in vertex-wise amygdala surface area across patient groups carrying a supernumerary X-chromosome (Pearson’s r for inter-vertex variation in area change ranging from 0.60 to 0.73), and far fewer significant differences in XYY. However, when examining the spatial distribution of t-statistics before FDR correction, surface area changes in XYY were similar to surface area changes in the other three karyotype groups of interest (Pearson’s r ranging from 0.6 to 0.72, Fig 2C). This result suggests that a supernumerary Y-chromosome exerts effects on amygdalar shape that are weaker than but nonetheless spatially convergent with effects of a supernumerary X-chromosome. The general pattern of shape changes suggests that the rostral surface of the amygdala is focally contracted in sex chromosome aneuploidies to a degree not expected based on total amygdala surface area.

### Hippocampal shape is systematically altered by SCA

We applied a similar analysis strategy to the hippocampus surface to assess whether hippocampus shape is atypical in sex chromosome aneuploidies. After FDR correction, we observed a similar pattern of significant surface area difference to our amygdala analyses (Fig. 2B). In particular, we identified significant differences across X aneuploidy groups relative to their respective gondal controls that were observable in attenuated form in the contrast between XYY and XY groups (Pearson’s r ranging from 0.70 to 0.83, Fig. 2C). This pattern included bands of focal contraction along the ventral and caudal surface of the hippocampus, and a patch of focal contraction on the rostral surface.

## Discussion

Our analyses provide several insights into the influences of sex and sex chromosome complement on anatomical organization of the amygdala and hippocampus, as detailed below.

### Amygdala and hippocampus volume and shape are not sexually dimorphic

Our findings replicate recent meta-analyses establishing that average amygdala and hippocampus volume are not statistically-significantly different between typically developing males and females after proper correction for sex-differences in total brain volume (Marwha, Halari, & Eliot, 2017; Tan, Ma, Vira, Marwha, & Eliot, 2016). In particular, although we observed large sex differences in raw amygdala and hippocampus volumes and an opposite effect in total-brain-volume-normalized hippocampal volume, these results were nonsignificant after both covariation- and allometry-based total brain volume correction. We conclude that gross differences in total brain volume between sexes, coupled to a hypoallometric scaling regime for the amygdala and hippocampus, give rise to false sex differences in raw and normalized region volumes. We also used high-resolution surface-based shape assays to demonstrate that amygdala and hippocampus shape are also not significantly different between males and females. These results are in contrast to a previous report of sex differences in amygdala shape assessed via a different method (Kim et al., 2012), suggesting that choice of processing pipeline may influence detection of significant sex differences. Future studies should systematically assess consequences of processing pipeline choice to explain these disparate findings, as has been done for other subcortical compartments (Makowski et al. 2018). In light of recent evidence for publication bias in neuroimaging of sex differences, our null findings are an important contribution to a body of literature that generally overestimates sex differences in the human brain (David et al., 2018). Thus, well-established sex-biases in the prevalence rates of psychiatric disorders that are linked to the amygdala and hippocampus do not appear to be accompanied by clear sex-biases in the macroanatomical organization of these structures in a typically developing population.

### Carriage of supernumerary sex chromosomes changes amygdala volume, but not hippocampus volume

We found that supernumerary sex chromosomes decrease the proportional volume of the amygdala. This decrease was common across aneuploidy groups, and mostly consistent across techniques to account for total brain size. The exception to this generally observed decrease was the XXX/XX contrast with normalization based correction for total brain volume, where no significant difference in amygdala volume was detected. Converging evidence across other total brain volume correction techniques and related patient groups suggests that this null finding may arise from methodological issues with normalization by division. Future simulation studies should assess why normalization by division yields different results, and in what situations it is or is not an appropriate technique to control for total brain volume. This amygdala volume decrease in patients carrying supernumerary sex chromosomes is consistent with a previously observed amygdala volume increase in patients with X-chromosome monosomy (Turner’s syndrome) (Kesler et al., 2004). These results suggest that amygdala volume scales negatively with mounting sex chromosome dosage, and indicate that the increases in both X- and Y-chromosome dosage can induce disproportionate reductions in amygdala volume relative to their effects on overall brain size. Although we detected differences in raw hippocampal volume in XXX females and XXY males relative to controls, these effects were not significant after total brain volume correction, suggesting that raw volume changes may be secondary to changes in total brain volume.

### Carriage of supernumerary sex chromosome alters amygdala and hippocampus shape

Carriage of additional X- and Y-chromosomes were both associated with a similar topography of shape change in the amygdala and hippocampus, although visual examination of FDR-corrected maps (Fig. 2B) show clearly that the extent of significant effects was greater in X-versus Y-chromosome aneuploidy. In particular, an area along the rostral surface of the amygdala centered on the basomedial nucleus was focally contracted in patients carrying supernumerary sex chromosomes. Our report of spatially specific contraction in X-chromosome supernumeracy, in combination with a previous report of spatially specific expansion of the basomedial nucleus in X-chromosome monosomy (Green et al., 2016), suggests a spatially specific X-chromosome-dosage effect on amygdala shape. In the hippocampus, we observed a large band of areal contraction along the midsection of the rostral-caudal axis, as well as a small area of focal expansion along the caudal surface. These areas of shape change cut across hippocampal subfields, suggesting that SCA does not target particular hippocampal compartments. Importantly, we show that SCA effects in amygdala and hippocampus shape are not a simple by-product of changes in the overall size of these structures. Future studies should confirm our findings with high-field MRI images, and align observed shape changes to known functional, connectivity, and cytoarchitectonic gradients within the amygdala and hippocampus. These studies would ideally identify core subregional modules of vulnerability to copy number variation that explain elevated rates of psychiatric disorders seen in SCA (Hong & Reiss, 2014).

## Conclusions

Carriage of supernumerary X- and Y-chromosomes exert convergent effects on the volume and shape of the amygdala and hippocampus. This convergence across the entire family of SCAs echoes convergent effects previously identified in the cerebral cortex (Raznahan et al., 2016), striatum and thalamus (Reardon et al., 2016), and cerebellum (Mankiw et al., 2017). This robust pattern suggests that SCA influence on brain anatomy arises from the shared gene content between X- and Y-chromosomes (i.e. pseudoautosomal regions, PAR). Indeed, recent *in-vitro* gene expression results have shown a selective enrichment for PAR gene expression in SCA patients (Raznahan et al., 2017).

These findings extend extant literature examining neuroanatomy in SCA patients in three key ways. First, our results are the first characterization of limbic anatomy changes in 47,XYY patients. The finding of reduced amygdala volume in these patients helps explain why Y-chromosome aneuploidy is often accompanied by social deficits and autistic symptoms (Lee et al., 2012), given well-documented patterns of amygdala changes in autistic patients (Avino et al., 2018; Nordahl et al., 2012). Secondly, the present findings highlight the power of a comparativist approach that profiles brain phenotypes across a family of related disorders, rather than in isolation; this approach has identified robust biomarkers across a range of brain disorders and imaging assays (Goodkind et al., 2015; Kaczkurkin et al., 2017; Spronk et al., 2018). This approach helps identify convergent effects across many genetic insults and reinforces reproducibility of findings across variation in genetic copy number. Thirdly, our results demonstrate how high-resolution surface analyses can complement traditional bulk volume analyses. In particular, although SCA was not accompanied by a change in hippocampal volume, our surface-based analysis identified a robust, spatially consistent change in hippocampus shape that may have consequences for the function of neuronal circuits therein.

The present study has three key limitations. We do not address behavioral correlates of the observed changes in limbic anatomy; performing such analyses would be a critical to fully assess whether limbic anatomy is a biomarker for phenotypic severity. Additionally, because our sample is cross-sectional, we cannot assess how observed effects are patterned across developmental time within individuals. Although longitudinal studies are difficult in rare patient populations, these data would help clarify the temporal properties of limbic anatomy development in SCA. Lastly, although structural MRI is a powerful tool for assessing gross regional volume and shape, it does not allow direct embedding of our present findings within known microarchitectural gradients of the limbic system (Janak & Tye, 2015).

Notwithstanding these limitations, our study is a major step towards understanding how copy number variation shapes the brain systems that underlie risk for psychopathology and, more broadly, social and emotional life. Understanding how a range of SCAs cause psychopathology via limbic brain systems may help triangulate stereotyped risk pathways that can serve as biomarkers and targets for intervention.

